# High-Frequency Activity Encodes the Temporal Dynamics of Hierarchical Prediction Errors in Humans: An Electrocorticography Study

**DOI:** 10.1101/2025.11.29.691270

**Authors:** Yiyuan Teresa Huang, Yuan-Yuan Chen, Shigeta Fujitani, Seijiro Shimada, Shinsuke Koike, Naoto Kunii, Nobuhito Saito, Zenas C. Chao

## Abstract

Prediction error refers to the discrepancy between expected and actual sensory input. Its hierarchical organization has been demonstrated through decomposed brain responses in the local-global paradigm, where temporal regularities are established locally at individual stimulus transitions and globally across multi-tone sequence structures. In macaques and marmosets, local and global prediction-error signals have both been found in high-frequency oscillations (60-150 Hz) recorded with ECoG. In humans, however, these signals have primarily been observed in a lower frequency range (30-100 Hz) using EEG. Here, we recorded human ECoG to achieve higher signal fidelity, enabling examination of neural oscillations above 100 Hz (high-frequency activity, HFA), which are believed to be closely linked to local spiking activity. Eight participants listened to auditory sequences that either followed their local and global regularities (local and global standards) or violated them (local and global deviants). Robust HFA responses were observed for the local deviants, but these responses were reduced when the deviants were expected based on the global regularity, indicating both levels of prediction-error processes contribute to the observed activity. Importantly, these HFA responses could be decomposed into two subcomponents: an early component reflecting local prediction-error signals localized to lateral auditory regions, and a late component reflecting global prediction-error signals prominent in both lateral auditory and frontal cortices. Together, these findings demonstrate that neural oscillations above 100 Hz encode hierarchical prediction errors not only in non-human primates but also in humans.

## Introduction

Predictive coding postulates that the brain constructs an internal model to anticipate upcoming sensory events, and updates this model when actual input deviates from those predictions through the generation of prediction errors (Clark, 2013; Friston, 2010; Rao & Ballard, 1999). A substantial body of research has demonstrated neural representations of prediction-error signals, such as mismatch negativity (MMN) evoked in a classical auditory oddball paradigm (Garrido et al., 2009; Näätänen et al., 2007). Importantly, prediction errors exhibit hierarchical organization (El Karoui et al., 2015; Goris et al., 2018; Sauer et al., 2017; Wacongne et al., 2011), a feature that has been successfully examined using the local-global oddball paradigm (Bekinschtein et al., 2009). In this paradigm (Figure 1), violations occur either at a local transition level or across the global sequence structure, enabling interaction of local and global prediction-error signals. Researchers have successfully extracted local and global prediction-error signals in non-human primate and human studies. For example, using electrocorticography (ECoG), which directly records neural activity from the cortical surface, studies in both macaques (Chao et al., 2018) and marmosets (Chao et al., 2024; Jiang et al., 2022) have reported that these two prediction-error signals appear in gamma oscillations extending into the high-frequency range (>100 Hz) and with distinct temporal delays. On the other hand, in humans, scalp EEG studies have similarly distinguished early and late gamma responses (<100 Hz) (Chao et al., 2022) and early and late MMN subcomponents (Huang et al., 2024) associated with local and global prediction errors.

**Figure 1.**
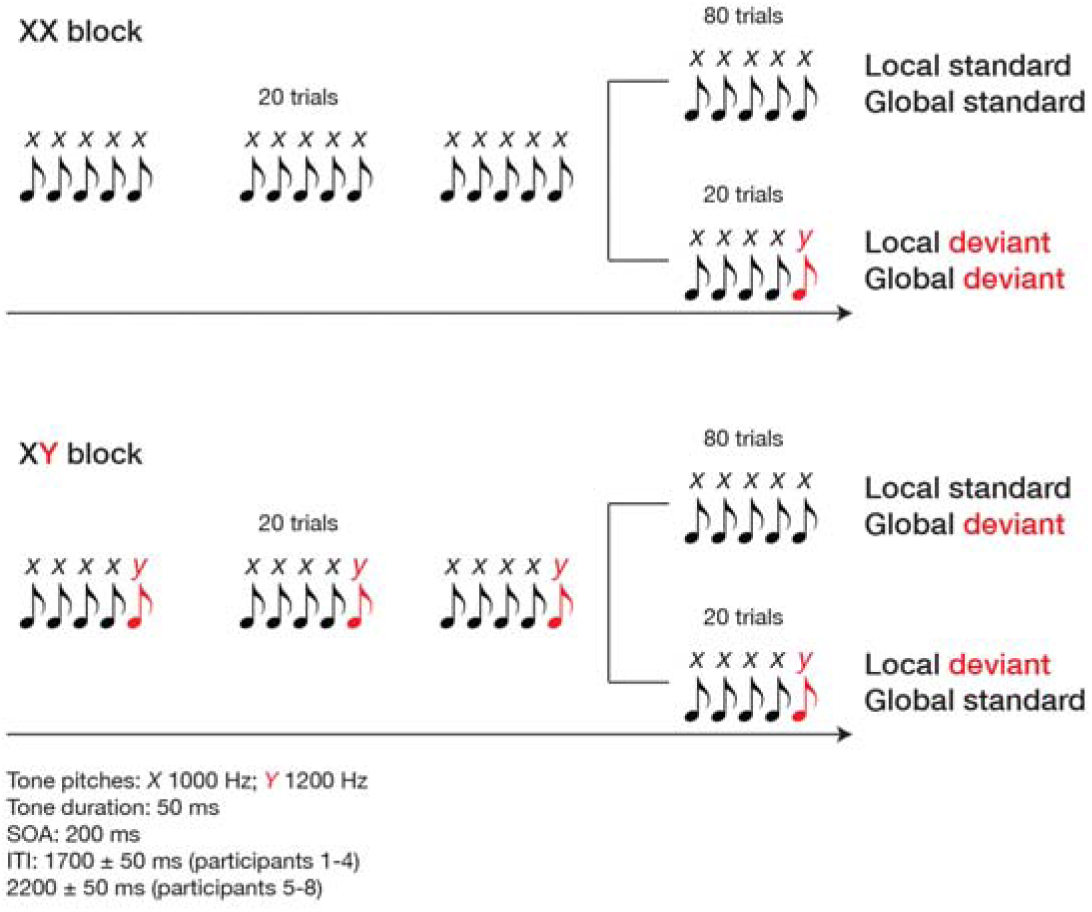
Task design. Abbreviations: SOA, stimulus onset asynchrony. ITI, inter-trial interval.

However, while human scalp EEG is a noninvasive and widely accessible recording system, its signals cannot reliably capture high-frequency activity (HFA; >100 Hz) (Muthukumaraswamy, 2013; Whitham et al., 2007) or provide precise localization of neural sources due to spatial blurring by the skull and scalp and volume conduction in the brain (Michel & Brunet, 2019). HFA has been proposed as an electrophysiological signature of local neuronal activity, reflecting either increased firing rates or enhanced neural synchrony within a small cortical area (Ray et al., 2008; Sinai et al., 2009). Moreover, this high-frequency response has been observed to emerge earlier for deviant stimuli compared with low-frequency event-related potentials (Dürschmid et al., 2016). Nevertheless, it remains unclear whether, using human ECoG with higher signal fidelity in both frequency and spatial domains, these two prediction-error signals are also represented in HFA with distinct temporal profiles that reflect hierarchical predictive processing, and how such representations are organized across different cortical regions. Verifying whether HFA encodes hierarchical prediction errors in humans, as observed in non-human primates, is essential for establishing the translational validity of hierarchical predictive coding. Demonstrating similar HFA correlates in human ECoG would strengthen the link between microcircuit mechanisms identified in animal studies and large-scale cortical computations underlying human perception and cognition.

In the present study, we use ECoG to record high-resolution neural activity from epilepsy patients while they perform the auditory local-global oddball paradigm. This paradigm combines local and global regularities with standard and deviant stimuli, resulting in four sequence types. We examine how oscillatory responses (4∼200 Hz) to these sequences varied across cortical regions, frequency bands, and time. Robust responses evoked by the local deviant compared with the local standard are observed in the high-frequency range (>100 Hz) and differ depending on whether the deviant was globally predictable (global standard) or less predictable (global deviant). The difference suggests that HFA reflects the mixture of local and global predictive processing. To dissociate these effects, we use parallel factor analysis (PARAFAC), previously applied to human EEG for analyzing prediction errors (Chao et al., 2022, Huang et al., 2024), to identify early and late subcomponents whose contributions correspond to local and global prediction-error signals, respectively. The early subcomponent is primarily localized to lateral auditory regions, whereas the late subcomponent is evident in both lateral auditory and frontal cortices. Together, these results indicate that HFA provides a shared neural substrate for prediction errors across species, with hierarchical interactions processed in a temporally ordered and spatially organized manner across the cortex.

## Materials & Methods

### Participants

Eight patients (4 males and 4 females; mean age ± standard deviation: 34.5 ± 9.7 years) with refractory epilepsy underwent subdural electrode implantation. The implantation sites were selected solely for clinical purposes, specifically to identify epileptic foci and to map critical functional areas surrounding the seizure focus (see Table 1). All participants met the inclusion criteria, which included self-reporting and being screened to confirm no significant auditory impairments, and no history of neurological or psychological disorders. The study was approved by the Research Ethics Committee of the Faculty of Medicine of the University of Tokyo (approval number: 1797), and written informed consent was obtained from all participants after they received a detailed explanation of the experimental procedures.

**Table 1.** Demographic information of participants.

|  | Age | Gender | Epileptic duration (yrs) | Epileptic foci | Electrode coverage | Total number of electrodes |
| --- | --- | --- | --- | --- | --- | --- |
| 1 | 39 | M | 30 | Lt mesial T | Lt FTP | 190 |
| 2 | 21 | F | 15 | Lt lat post T | Lt FT | 50 |
| 3 | 44 | F | 20 | Lt mesial T | Lt FTP | 190 |
| 4 | 45 | M | 26 | Bil. mesial T | Bil. FTP | 140 |
| 5 | 44 | M | 26 | Rt OF | Bil. FT | 92 |
| 6 | 32 | F | 12 | Rt OF, Rt mesial T | Lt FT, Rt FTP | 230 |
| 7 | 25 | F | 17 | Rt mesial T, Rt posterior CG | Lt O, Rt FTPO | 58 |
| 8 | 26 | M | 11 | Rt mesial T | Lt T, Rt FT | 208 |

Lt: left hemisphere; Rt: right hemisphere; F: frontal; T: temporal; P: parietal; O: occipital; OF: orbitofrontal; CG: cingulate gyrus

### Electrode placement and data acquisition

Subdural platinum electrodes (Unique Medical, Tokyo, Japan) were arranged in grids with silicone sheets. Two grid types were employed: one with 3-mm-diameter electrodes spaced 10 mm apart (center-to-center), and the other with 1.5-mm-diameter electrodes spaced 5 mm apart. The choice of the grid type and implantation sites was determined by the clinical needs of each participant. Reference electrodes were placed on the inner surface of the skull, facing the dura mater. Excluding the reference electrodes, a total of 1,158 electrodes were implanted across all participants: 644 in eight left hemispheres and 514 in five right hemispheres (see Table 1 for detailed location information).

The duration of electrode implantation for each participant typically ranged from 2 to 4 weeks. ECoG data were recorded using a multichannel EEG system (EEG 1200; Nihon Kohden, Tokyo, Japan) with a sampling rate of 2000 Hz and a band-pass filter of 0.09–600 Hz. The data were online-referenced to the reference electrodes. The data acquisition was conducted only after sufficient seizure information had been obtained for clinical purposes. No epileptic seizures were observed within 24 hours before or after the experiment.

### Auditory local-global oddball paradigm

In this paradigm (Figure 1), tone *x* was delivered as a low-pitched tone, and tone *y* as a high-pitched tone (1000 Hz vs. 1200 Hz for participants 1–2; 595 Hz vs. 1000 Hz for the remaining participants). Two types of five-tone sequences were used: one consisting of five low-pitched tones (*xxxxx*), and the other consisting of four low-pitched tones followed by a high-pitched tone (*xxxxy*). Each tone had a duration of 50 ms, with an stimulus onset asynchrony (SOA) of 200 ms. Two blocks were used, each consisting of 120 tone sequences (trials). In the XX block, 100 trials of the *xxxxx* sequence and 20 trials of the *xxxxy* sequence were presented. In the XY block, the proportions were reversed, with 20 *xxxxx* trials and 100 *xxxxy* trials. The inter-sequence onset interval was set at 1700 ± 50 ms for participants 1–4, and 2200 ± 50 ms for the remaining participants. The sequence order within each block was pseudo-randomized. 20 trials of the frequent sequence for that block (e.g., *xxxxx* in the XX block) were presented first, followed by a randomized order of the remaining 100 trials (e.g., 80 *xxxxx* and 20 *xxxxy* in the XX block). Each block was presented once. During each block, participants were instructed to attend to the sounds and count the number of infrequent tone sequences.

### Data analysis

#### Preprocessing

The preprocessing was conducted using MATLAB-based toolboxes: EEGLAB and FieldTrip. The raw data were downsampled to 500 Hz (function: *pop_resample.m*) and processed with a high-pass filter at 0.1 Hz and a notch filter at 50 Hz (functions: *pop_eegfiltnew.m* and *ft_preprocessing.m*). Epochs were then extracted from –1.4 s to 1.05 s relative to the onset of the fifth tone in each sequence (function: *pop_epoch.m*). We manually removed bad epochs and electrodes that exhibited excessive fluctuations, high-frequency or high-amplitude noise, or epileptic spikes. In total, 164 electrodes and 349 epochs (out of 1920) were excluded from further analysis (see Table 1). A common average reference method was done for each cortical grid for each participant (function: *ft_preprocessing.m*).

#### Event-Related Spectral Perturbation (ERSP)

For each participant, electrode, and trial, the time-frequency representation (TFR) of the processed signal was computed using a Morlet wavelet transformation with 200 center frequencies ranging from 1 to 200 Hz and a fixed wavelet cycle length of 7 (function: *ft_freqanalysis.m*). To extract event-related spectral perturbations, baseline normalization was applied using the decibel method, with the baseline period defined as –1 to –0.8 s relative to the fifth tone onset (i.e., –0.2 to 0 s relative to the first tone onset) (function: *ft_freqbaseline.m*). The resulting ERSPs were then averaged across trials for each sequence type within each block, for each participant and electrode.

#### Significant contrast responses

To quantify mismatch responses in the local-global oddball paradigm, we compared ERSPs from sequence *xxxxy* (local deviant) with those from sequence *xxxxx* (local standard), calculated as *xxxxy* – *xxxxx*. Importantly, this contrast was computed in both blocks, including the condition where *xxxxy* served as the global standard. This approach follows previous studies examining MMN elicited by local violations under varying levels of global expectation (Chao et al., 2022; Goris et al., 2018; Huang et al., 2024; Sauer et al., 2017; Wacongne et al., 2011). We then tested whether the contrast response at each electrode significantly differed from zero within each participant. Specifically, for each participant, the *xxxxy* dataset was compared with the *xxxxx* dataset. Both datasets were structured in three dimensions: trials, time points, and frequency bands. To reduce computational load, the analysis was restricted to the time window from 0 to 0.6 seconds relative to the onset of the fifth tone. Frequency resolution was further adjusted in 4-40 Hz in 2-Hz increments; 44-100 Hz in 4-Hz increments; 108-200 Hz in 8-Hz increments, which reduced the total number of frequency points from 200 to 46. Statistical testing was conducted using an independent-samples t-test with the Monte Carlo method and cluster-based correction across time and frequency domains (function: *ft_freqstatistics.m*). The analysis included 500 randomizations with an alpha level of 0.05.

Figure 2A shows examples of ERSPs for the two sequences across the two blocks, along with their contrast responses (significant responses are outlined in black) from a right temporal grid electrode in participant 6.

**Figure 2.**
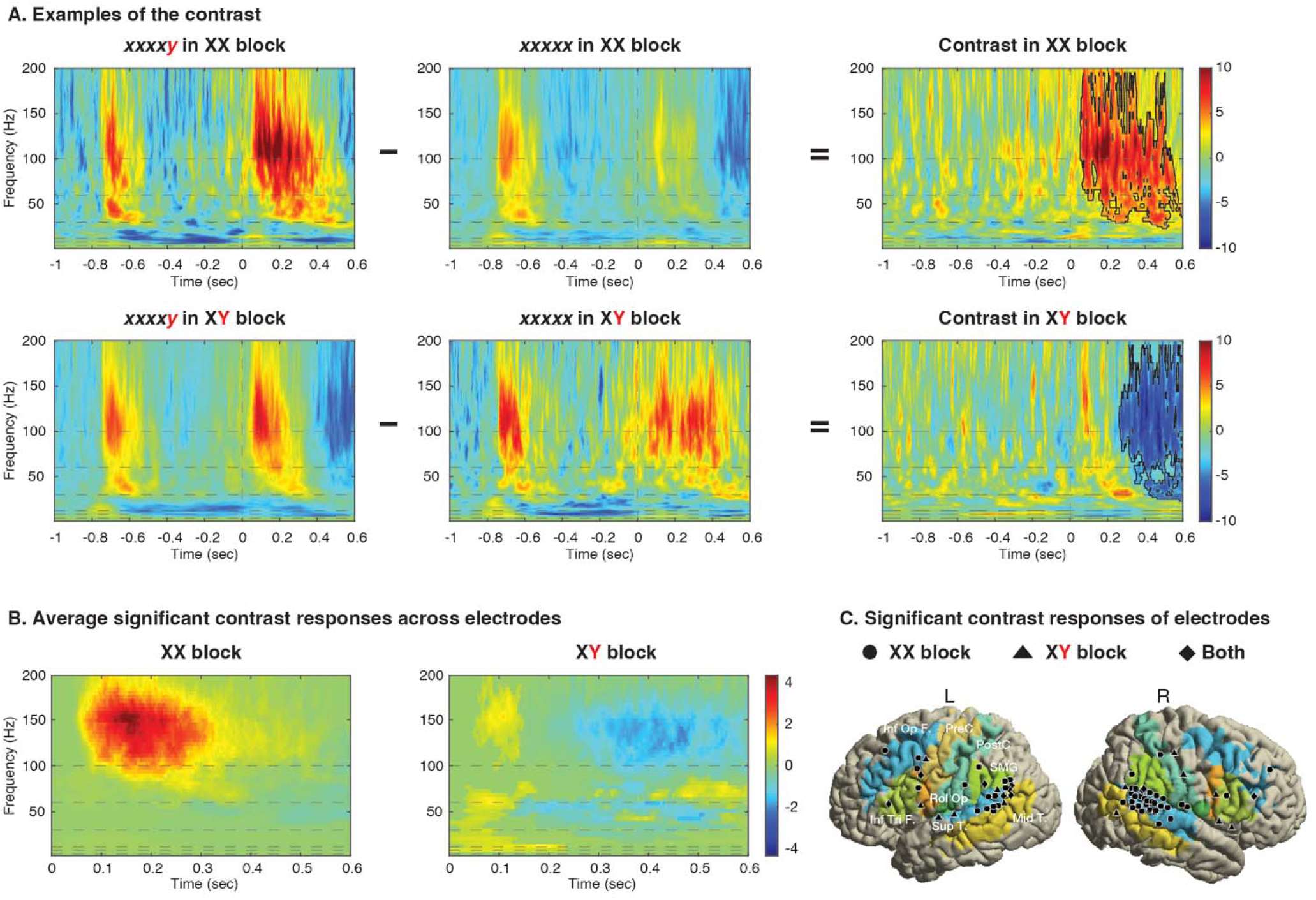
Event-related spectral perturbations (ERSPs) and significant contrast responses. A. Example ERSPs for the four sequence types and their contrasts from a right temporal grid electrode in participant 6. The vertical line indicates the onset of the last stimulus in the sequence. Horizontal lines mark frequencies at 4, 8, 12, 30, 60, and 100 Hz. B. Significant contrast responses after the last stimulus, obtained by averaging contrast responses across all electrodes, with non-significant values replaced by zero. C. Electrodes showing significant contrast responses for the XX-only, XY-only, and both-block conditions, marked with circles, triangles, and diamonds, respectively, on the cortical surface. Abbreviations: Inf Op F., inferior opercular frontal; Inf Tri F., inferior triangular frontal; PreC, precuneus; PostC, postcentral; Rol Op, Rolandic operculum; Sup T., superior temporal; SMG, supramarginal gyrus; Mid T., middle temporal.

### Electrode localization and visualization

To visualize electrodes on cortical surfaces, we followed three main procedures: electrode localization, cortical surface extraction, and re-alignment of electrode locations. First, post-operative computed tomography (CT) images were registered to pre-operative magnetic resonance imaging (MRI) scans using a mutual information method in Dr. View/Linux (Infocom, Tokyo, Japan). Subdural electrodes were then reconstructed and visualized on each participant’s three-dimensional (3D) brain surface using Real INTAGE (Cybernet Systems, Tokyo, Japan). Electrode coordinates were subsequently transformed into Montreal Neurological Institute (MNI) space using the normalization method in SPM8 (Wellcome Department of Imaging Neuroscience, London, United Kingdom). Second, cortical surfaces were extracted from each participant’s structural MRI using FreeSurfer. Although the post-operative CT had been aligned to the pre-operative MRI, electrode locations could still be slightly misplaced on the cortical surface. To correct this, we applied the re-alignment method described by Dykstra et al. (2012) (Dykstra et al., 2012), which projects each electrode grid onto the cortical hull (function: *ft_electroderealign.m*).

After electrodes were first realigned to each participant’s cortical hull, we repeated the procedure with a standardized cortical surface (the MNI average of 152 T1-weighted MRIs) to enable group-level visualization and comparison. Using the processed electrode placements and the standardized surface, electrode locations can be visualized using the functions: *ft_plot_mesh.m*, *ft_plot_sens.m*. To map electrode locations onto brain regions, we used the Automated Anatomical Labeling atlas 3 (AAL3) with the function *ft_plot_mesh.m*.

### Parallel factor analysis

To extract underlying subcomponents from the HFA contrasts across the dimensions of two blocks * time points * two regions of interest, we applied parallel factor analysis (PARAFAC)—a generalization of principal component analysis to higher-order arrays (Harshman & Lundy, 1994). The analysis was performed using the N-way Toolbox (Andersson & Bro, 2000) (function: *parafac.m*), with the number of target subcomponents set to two. The resulting subcomponents were visualized in their original dimensions of Block, Time course, and Brain region. The first dimension describes how strongly the HFA contrast in each block is contributed to by each subcomponent, and each subcomponent consists of temporal and spatial activation profiles (the second and third dimensions).

To verify the functional roles of the two subcomponents, we employed a quantitative predictive-coding model that estimates the size of local and global prediction errors in the two blocks of the local–global oddball paradigm (Chao et al., 2022). Using the published code, we obtained model-derived values of local and global prediction errors. The local prediction-error values were positive in both block contrasts, whereas the global prediction-error values were positive in the XX block and negative in the XY block. These value patterns were used to determine the functional correspondence of the two subcomponents.

## Results

### Prediction errors in high-frequency activity predominantly in auditory areas

For each block, we extracted prediction-error responses by contrasting ERSPs between sequences *xxxxy* and *xxxxx* (*xxxxy* – *xxxxx*). Figure 2A shows an example of ERSPs from a right temporal grid electrode in participant 6. In this example, significant contrast responses are outlined in black (*t* = 1.94e^4^, *p* = 0.002 in the XX block; *t* = –1.23e^4^, *p* = 0.002 in the XY block) (independent-samples t-test, Monte Carlo method with 500 randomizations, α = 0.05, two-tailed, cluster-based correction). The contrast responses in the XX block show positive significant differences in a broad frequency range from 30 to 200 Hz, appearing immediately after the last tone onset. On the other hand, those in the XY block show negative differences in a similar frequency range but occurring around 0.3 s.

To characterize the overall significant contrast responses, Figure 2B presents the responses averaged across all electrodes and participants for both blocks, after replacing non-significant values with NaN. In the XX block, a positive difference was observed mainly in the high-frequency band (100–200 Hz) between 0.1 and 0.3 s after the onset of the last tone. In the XY block, weak positive differences appeared in the theta, alpha, and beta bands (< 60 Hz) between 0 and 0.2 s, as well as in the high-frequency band around 0.1 s. These were followed by a strong negative difference in the high-frequency band (0.3–0.5 s) and a positive difference in the frequency range between 60 and 80 Hz. Overall, the HFA was evident in both blocks.

Figure 2C shows a total of 70 electrodes with significant contrast responses: 47 significant electrodes observed in the XX block only (circle), 13 in the XY block only (triangle), and 10 in both (diamond), with the surface colored according to the AAL atlas. Table 2 lists the number of significant electrodes per brain area and block. The majority were located in auditory regions (57 in total), including the right superior temporal (22 electrodes), left superior temporal (8), left middle temporal (7), and right supramarginal (5) areas. Additionally, 13 significant electrodes were observed in frontal regions. These findings indicate that prediction-error responses during the local-global oddball paradigm were localized to both sensory-related and frontal areas, consistent with the previous intracranial study of El Karoui et al. (2015). Supplementary Figure 1 further shows averaged contrast responses in significant electrodes and frequency bands of HFA across the defined time windows.

**Table 2.** Significant electrode locations based on AAL atlas.

|  | XX block | XY block | Both | Subtotal |
| --- | --- | --- | --- | --- |
| Left middle frontal | 3 |  |  |  |
| Right middle frontal | 3 |  |  |  |
| Left inferior triangular frontal |  | 1 | 1 |  |
| Right inferior triangular frontal | 1 |  |  |  |
| Left inferior operculum frontal | 1 |  | 1 |  |
| Right inferior operculum frontal | 2 |  |  | 13 |
| Left rolandic operculum |  |  | 1 |  |
| Right rolandic operculum |  | 1 |  |  |
| Left precentral | 1 |  |  |  |
| Left postcentral |  | 1 |  |  |
| Right postcentral | 1 | 3 |  |  |
| Left supramarginal | 3 | 1 |  |  |
| Right supramarginal | 2 | 2 | 1 |  |
| Left superior temporal | 4 | 1 | 3 |  |
| Right superior temporal | 20 | 2 |  |  |
| Left middle temporal | 4 |  | 3 |  |
| Right middle temporal | 2 | 1 |  | 57 |
| Total | 47 | 13 | 10 | 70 |

Two key observations can be found in these results. Overall, the number of significant electrodes is roughly balanced between the two hemispheres. However, a closer look reveals that prediction-error contrast responses in the XX block were predominantly observed in the right superior temporal area (20 electrodes), compared with the left (5 electrodes). One possible reason is that participant 6 contributed many significant electrodes (10) but had coverage only in the right temporal region. To explore potential hemispheric lateralization, we examined participant 4, the only case with bilateral temporal coverage. In this participant, four electrodes on the right side and none on the left showed significant responses. While this hints at possible right-hemisphere dominance, conclusions remain tentative given the limited sample size. Future studies with broader bilateral coverage are needed to verify hemispheric lateralization in prediction-error processing.

Second, prediction-error responses were smaller in the XY block than in the XX block, both in response magnitude (Figure 2B) and number of significant electrodes (Figure 2C and Table 2). This reduction aligns with prior findings showing that MMN amplitude decreases when a local deviant appears within a frequent multi-stimulus sequence (i.e., *xxxxy* in the XY block) compared to an infrequent one (*xxxxy* in the XX block) (Goris et al., 2018; Huang et al., 2024; Sauer et al., 2017). Moreover, the attenuation of HFA observed in our study also suggests that this activity comprises multiple subcomponents. Specifically, due to the combination of local/global and standard/deviant sequences, their deviant effects vary, resulting in different overall HFA amplitudes. To test this hypothesis, we performed PARAFAC on the two contrast responses within each region of interest.

### Early and late subcomponents for local and global prediction errors

We focused on the ERSP contrasts in the high-frequency band, averaged separately across significant electrodes in the lateral auditory cortex (including superior and middle temporal areas) and frontal areas (Figure 3A). In the XX block, the auditory HFA contrast (red solid line) shows a marked increase from 0.1 to 0.3 s after the final tone, whereas in the XY block (red dashed line), it exhibits only a small initial rise followed by a mild sustained decrease. The frontal HFA contrasts (blue solid and dashed lines) display similar temporal patterns as those in the auditory region, but with smaller power.

**Figure 3.**
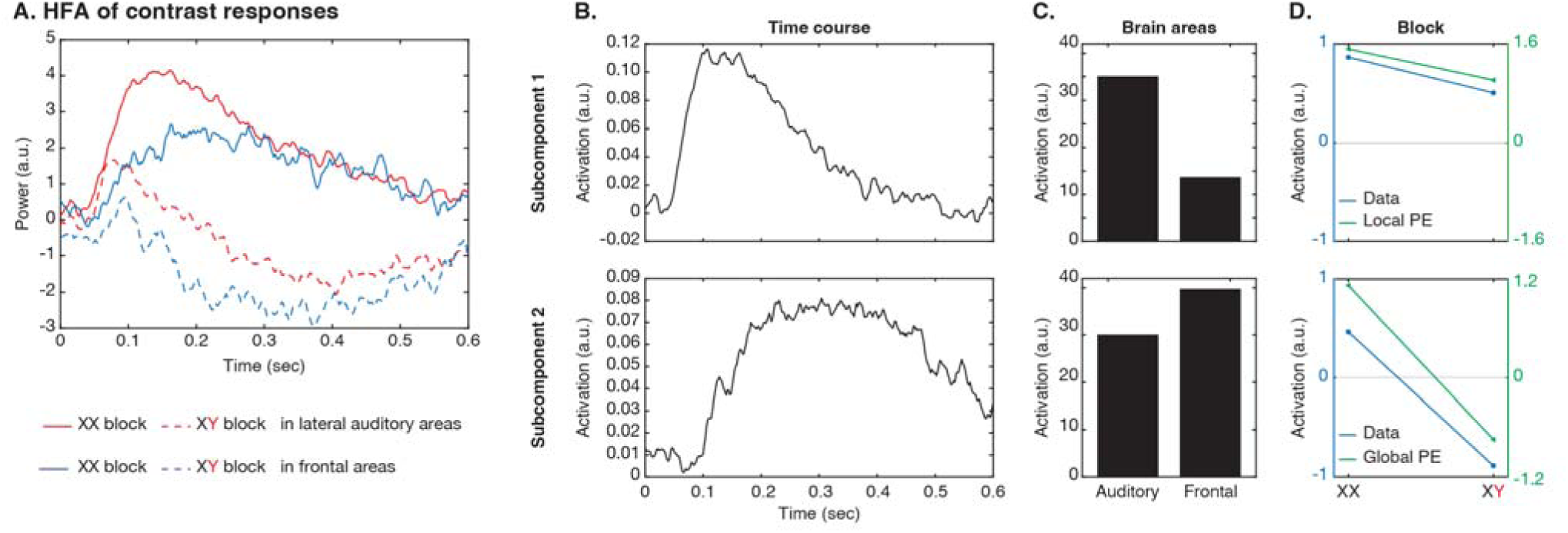
High-frequency activity (HFA) of contrast responses and neural representations of local and global prediction-error signals. A. Four contrast response curves for the two blocks, obtained by averaging significant electrodes within two cortical areas. The four curves were decomposed into two subcomponents, and subcomponent activations are shown in (B) Time course, (C) Brain areas, and (D) Block dimensions. In panel D, the block dimension reflects the activation of each subcomponent (blue line), while the model-derived values for the local and global prediction errors are shown in green.

To demonstrate that HFA reflects hierarchical prediction-error processes, we performed PARAFAC on these two contrast responses across the two regions of interest (a total of four curves). Two HFA subcomponents were extracted, jointly explaining 100% of the variance in the dataset. The first subcomponent peaked at 108 ms (Figure 3B) and was predominantly contributed by the lateral auditory cortex (Figure 3C). The second subcomponent increased in the time period between 0.2 and 0.5 s (peaking at 304 ms) in both frontal and auditory regions. To interpret their functional significance, we compared the subcomponent contributions (Figure 3D, blue line) with model-derived values of local and global prediction errors (green line). The early HFA subcomponent shows positive contributions in both blocks, consistent with local prediction-error values. On the other hand, the late subcomponent displays a positive contribution in the XX block and a negative contribution in the XY block, corresponding to the global prediction-error values. The negative contribution in the XY block reflects the comparison between sequences *xxxxy* and *xxxxx* (global standard – global deviant). In sum, these results demonstrate that PARAFAC effectively dissociates subcomponents underlying HFA, indicating that local and global prediction errors are represented in temporally distinct HFA and spatially organized across cortical regions. Importantly, we confirmed that HFA reflects hierarchical predictive processing across species, extending from non-human primates to humans.

## Discussion

This study advances our understanding of the presence, functional significance, and cortical localization of hierarchical prediction error signals in the high-frequency band of the human brain. First, we combined the local-global oddball paradigm with human ECoG recordings to probe prediction-error signals at two hierarchical levels. We demonstrated robust deviant responses in the high-frequency range and further decomposed them into two subcomponents. The local prediction-error subcomponent was represented in early HFA, predominantly in the lateral auditory cortex, whereas the global prediction-error subcomponent was represented in late HFA in both auditory and frontal regions. In sum, this study not only reveals prediction-error signals extending into the high-frequency range but also highlights the composite nature of HFA, indicating hierarchical prediction-error processing, previously observed in non-human primates.

### Functions of high-frequency activity as prediction errors

Neural oscillations reflect the synchronized activity of neuronal populations, and such synchronization can strengthen or weaken connections between neurons. In the predictive coding framework, strong gamma oscillatory power reflects bottom-up transmission of mismatched sensory information to higher-order cortical areas (Bastos et al., 2012; Chao et al., 2022), followed by desynchronization in the alpha/beta bands that signals a “breaking” of existing predictions and allows new evidence to reshape internal models (Arnal & Giraud, 2012; Chao et al., 2018; Jiang et al., 2022). Intracranial electrophysiological recordings have shown that gamma activity is not confined to the 30–100 Hz range (low gamma) but can extend into frequencies above 100 Hz (high gamma) (Crone et al., 2006; Lachaux et al., 2012), and the low-and high-gamma bands are thought to reflect distinct neural mechanisms (Crone et al., 1998, 2001), although their exact frequency boundaries may vary across studies. In addition, when oscillatory responses are not limited to a specific frequency range due to recording spec but appear simultaneously across a broad spectrum, this broadband power is thought to represent the cumulative spiking activity of largely unsynchronized local neuronal populations (Jacobs et al., 2010). However, recent evidence suggests that increases in power between 60 and 200 Hz associated with attention may preferentially reflect network-level synchronization rather than a purely local rise in firing rate (Ray et al., 2008). Despite this apparent discrepancy, it is generally accepted that HFA serves as a reliable indicator of spatially localized cortical processing. In addition, the activity has been consistently observed when unpredictable or deviant stimuli are presented, and studies using many-standard control designs have shown that HFA (70–150 Hz) in the lateral superior temporal gyrus is specifically evoked by deviant detection rather than by sensory adaptation (Ishishita et al., 2019).

Beyond prediction errors elicited by a single violated regularity, Bekinschtein and colleagues introduced the local–global oddball paradigm to investigate prediction errors at both the local (tone-to-tone) and global (multi-tone sequence) levels. Our human ECoG study adopted this paradigm, building on earlier work by El Karoui and colleagues, who reported high-gamma contrast responses (*xxxxy* – *xxxxx*) across merged blocks in the 60–120 Hz range. We argue, however, that this contrast approach primarily captures the local prediction-error signal and overlooks the contribution of the global prediction error. From the perspective of the quantitative predictive-coding model (Chao et al., 2022), the sizes of local and global prediction errors calculated based on tone transition and sequence probabilities are different between blocks (i.e., local = 1.52, 1.12; global = 1.01, –0.75 for XX and XY blocks, respectively). Thus, the merged contrast response still contains overlapping contributions from both levels of prediction error (local = 1.52+1.12 and global = 1.01+[–0.75]).

To address this issue, we applied PARAFAC to the dominant HFA contrast responses in each block, enabling us to disentangle the underlying subcomponents. The analysis revealed two distinct HFA components: an early component (∼108 ms) primarily in the lateral auditory cortex representing local prediction errors, and a late component (∼304 ms) observed in both auditory and frontal regions representing global prediction errors. For comparison, our previous studies reported human MMN subcomponents peaking at 136 ms and 200 ms (Huang et al., 2024) and gamma subcomponents peaking at 156 ms and 192 ms (Chao et al., 2022). Although these HFA components did not generally peak earlier, they emerged around or even before 100 ms, highlighting the rapid dynamics of high-frequency responses that allow for quick prediction updating. In addition, the ECoG recordings provided precise spatial localization of hierarchical prediction-error signals, showing that short-timescale information is predominantly processed in sensory-related regions, whereas long-timescale information recruits additional frontal involvement. Moreover, our human findings echo results from monkey and marmoset studies, which similarly identified early subcomponents confined to temporal sites and late subcomponents spanning both temporal and frontal areas (Chao et al., 2018; Jiang et al., 2022).

Although our findings extend prediction-error representations into the high-frequency range, we did not observe strong modulation in the low-gamma band (30–100 Hz), which has been frequently reported in human EEG and in early ECoG studies using the local–global oddball paradigm. One likely explanation for this discrepancy is the number of trials. The HFA likely reflects spiking-related activity that can be observed even at the single-trial level, whereas low-gamma power may require a larger number of trials to be reliably detected in averaged data (Crone et al., 2001). Because the comfort and safety of postoperative patients were our top priorities, we limited task duration, resulting in fewer trials per condition. Nevertheless, the robustness of HFA despite fewer trials underscores its reliability as an index of hierarchical prediction-error signaling, even within the constraints of a busy clinical schedule. Importantly, these neural signatures in the local–global oddball paradigm could serve as neurophysiological markers for assessing predictive-processing capacity in clinical populations. Previous studies have shown that attentional allocation influences local and global regularity learning, and that atypical hierarchical prediction-error processing characterizes individuals with schizophrenia and autism spectrum disorder.

### Limitations and future research

Several limitations should be considered when interpreting these findings. First, As observed in the study, the seemingly larger number of significant electrodes in the right hemisphere—while acknowledging the uneven electrode distribution between hemispheres— may indicate (1) right-hemisphere dominance in spectral-feature processing for predicting tone frequency (Rinne et al., 2005) and (2) a possible predominance of mismatch-response generation in right frontal regions (Doeller et al., 2003; Opitz et al., 2002). However, because electrode placement in clinical ECoG is determined primarily by medical necessity, cortical coverage was uneven across hemispheres and regions, precluding a systematic evaluation of hemispheric lateralization. Future research combining datasets across patients could help clarify whether hierarchical prediction-error representations are lateralized or symmetrically distributed across the cortex. In addition, recent findings suggest that the detection of local and global regularity violations may occur at the subcortical level (Mazancieux et al., 2023). Future intracranial studies with greater access to subcortical structures will be crucial for providing direct evidence of where deviant detection initially arises in the human brain.

Second, our analyses were based on averaged responses across trials, which capture stable prediction-error components but not their trial-by-trial dynamics. Recent work suggests that prediction errors and subsequent prediction updates fluctuate dynamically across trials (Iglesias et al., 2013; Sedley et al., 2016). Applying single-trial computational modeling—such as Bayesian observer models or hierarchical Gaussian filters—could reveal how hierarchical prediction errors and updates evolve over time and across cortical areas.

Finally, although PARAFAC effectively dissociated temporal and spatial subcomponents, future studies should aim to link these subcomponents more directly to canonical predictive-coding circuits by integrating non-invasive neuroimaging, laminar recordings, and computational modeling. Moreover, examining how the early and late HFA subcomponents interact along bottom-up and top-down pathways will be essential for understanding the neural mechanisms underlying hierarchical predictive coding.

## Data availability

The raw anonymized data, as well as the data underlying all figures and supplementary figures, will be publicly available in the data repository upon publication.

## Author contributions

Y.T.H and Z.C.C. conceptualized the study. S.F., and N.K collected the data. Y.T.H and Y-.Y.C performed data analysis. Y.T.H wrote the paper, and Z.C.C., Y-.Y.C., S.F., S.S., S.K., N.K., and N.S edited the paper. All the authors contributed to and approved the final paper.

## Acknowledgements

This study is supported by the Japan Agency for Medical Research and Development (grant number JP19dm0207069) (to N.K.), and the World Premier International Research Center Initiative (WPI), MEXT, Japan (to Z.C.C.).

