## Supplementary Figure1 for "High-Frequency Activity Encodes the Temporal Dynamics of Hierarchical Prediction Errors in Humans: An Electrocorticography Study"

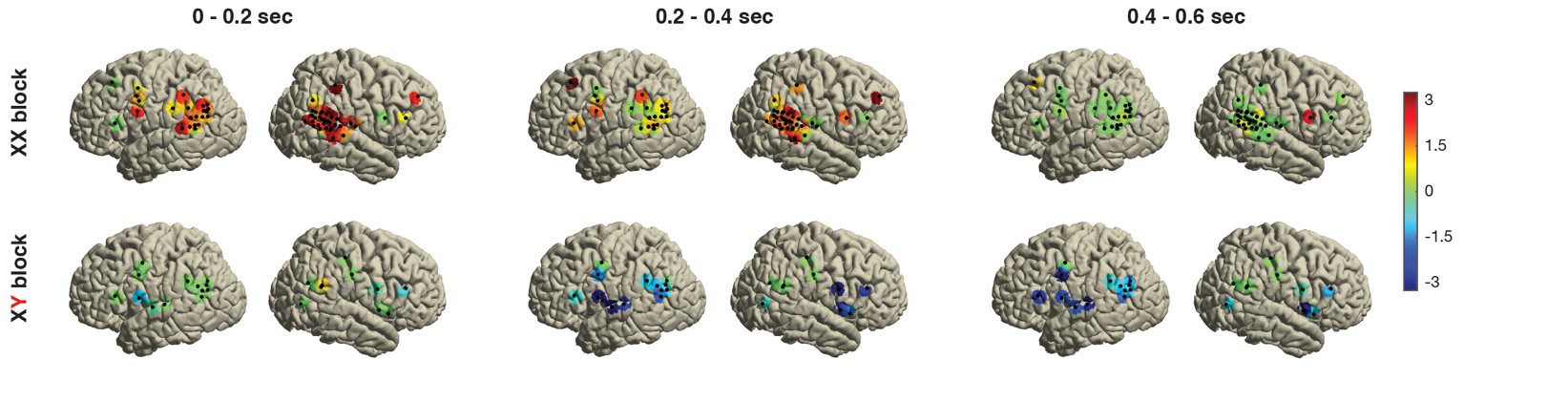

**Supplementary Figure 1.** Averaged contrast responses across significant electrodes and frequency bands above 100 Hz within the defined time windows.
